# Microfluidic deep mutational scanning of the human executioner caspases reveals differences in structure and regulation

**DOI:** 10.1101/2021.06.08.447609

**Authors:** Hridindu Roychowdury, Philip A. Romero

**Affiliations:** Department of Biochemistry, University of Wisconsin--Madison, Madison, WI, USA; Department of Chemical & Biological Engineering, University of Wisconsin--Madison, Madison, WI, USA; The University of Wisconsin Carbone Cancer Center, Madison, WI, USA

## Abstract

The human caspase family comprises 12 cysteine proteases that are centrally involved in cell death and inflammation responses. The members of this family have conserved sequences and structures, highly similar enzymatic activities and substrate preferences, and overlapping physiological roles. In this paper, we present a deep mutational scan of the executioner caspases CASP3 and CASP7 to dissect differences in their structure, function, and regulation. Our approach leverages high-throughput microfluidic screening to analyze hundreds of thousands of caspase variants in tightly controlled *in vitro* reactions. The resulting data provides a large-scale and unbiased view of the impact of amino acid substitutions on the proteolytic activity of CASP3 and CASP7. We use this data to pinpoint key functional differences between CASP3 and CASP7, including a secondary internal cleavage site, CASP7 Q196 that is not present in CASP3. Our results will open avenues for inquiry in caspase function and regulation that could potentially inform the development of future caspase-specific therapeutics.

## Introduction

Caspases are a ubiquitous family of cysteine proteases that play fundamental roles in programmed cell death and inflammation^1^. These enzymes have numerous ancillary roles in organismal development and homeostasis including cell differentiation, synaptic pruning, and cytokine processing^2,3^. In humans, there are twelve expressed members of the family, with Caspase-3, −6, −7, −8, and −9 primarily involved in apoptosis, and the others involved in pyroptosis and inflammation^1^. All caspases have a conserved core proteolytic domain and a variable N-terminal domain that’s involved with regulation of enzyme activity^3^.

Dysregulation of caspase activity is associated with cancer, neurodegeneration, vascular ischemia, and inflammatory diseases^1,4,5^. Consequently, these enzymes represent important therapeutic targets to treat a variety of human diseases^4^. However, despite their central role in human biology and disease, every caspase-targeting drug candidate has failed to pass through clinical trials^6,7^. A key challenge for therapeutic development has been the caspase family’s highly conserved proteolytic domain, which makes it difficult to selectively target one particular member and leads to off-target effects^8^. A deeper understanding of caspase structure, function, and regulation may eventually lead to small molecule modulators that selectively target members of the caspase family and open the door for novel therapeutics^6,9,10^.

In this work, we develop a high-throughput microfluidic platform for caspase screening and apply it to systematically map sequence-function relationships in the human executioner caspases^11,12^. Our microfluidic system consists of a fully integrated lab-on-a-chip that combines the addition of a fluorogenic substrate, incubation of the enzyme reaction, and fluorescence measurement. Our microfluidic chip can perform kinetics-based screening on millions of caspase variants. We applied our screening system to perform deep mutational scanning (DMS) on caspase-3 (CASP3) and caspase-7 (CASP7). The DMS data displayed known and expected signatures of caspase structure and function, but also revealed important differences between CASP3 and CASP7 that may be related to allosteric regulation and protein stability. Future exploration of the differences between human caspases may lead to more targeted drug design efforts.

## Results

### A microfluidic platform for ultra-high-throughput screening of caspases

High-throughput screening is an important tool for studying protein structure and function^13^. Caspases are challenging to screen because their activity cannot be readily linked cell growth or cellular fluorescence. Furthermore, caspases’ high catalytic rates make any cell-based assay difficult because the proteolytic cleavage reactions occur on significantly faster timescales than cell growth or fluorescent protein production. We developed a droplet microfluidic platform capable of *in vitro*, kinetics-based screening of millions of caspase variants.

Our microfluidic system encapsulates single *E. coli* cells, each expressing a unique caspase variant, into ~10 picoliter microdroplets that contain cell lysis reagents and a fluorogenic peptide substrate (Fig 1a). The droplets physically separate each cell and allow enzyme reactions to proceed in isolation. After encapsulation, the cells quickly lyse, releasing the expressed caspase and allowing it to interact with the substrate. The droplets are then incubated in an on-chip continuous flow reactor for ~3 minutes to allow the reaction to proceed. We found these short incubation times were necessary to separate highly active caspases from variants with severely attenuated activity. After incubation, each droplet is scanned with a laser fluorimeter and droplets displaying high fluorescence signals are sorted for downstream analysis. Our microfluidic platform is capable of screening 360,000 caspase variants per hour, while consuming only ~100 μL of assay reagents.

**Figure 1:**
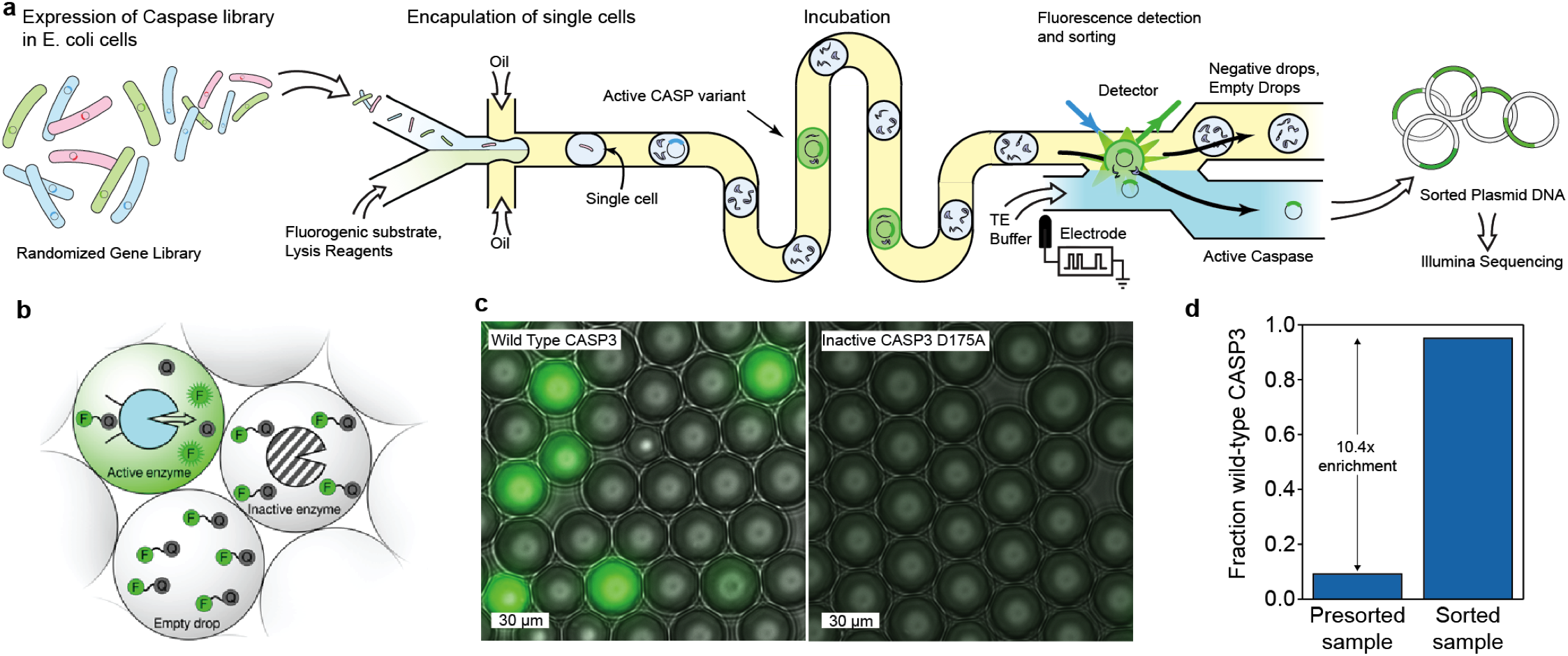
A droplet microfluidic platform for screening caspases. (a) A schematic of our microfluidic screening system. A dilute suspension of *E. coli* expressing caspase variants are injected into a microfluidic device and individual cells are encapsulated into microdroplets containing lysis reagents and a fluorogenic caspase substrate. The cells are lysed, the enzyme reaction is incubated on-chip, and the fluorescence of each droplet is analyzed using a laser. The fluorescent droplets are then sorted by electrocoalescence with an aqueous stream that collects the sorted plasmids for downstream analysis. (b) Droplets containing active caspase variants will fluoresce, whereas empty droplets and droplets containing inactive caspases will not. (c) Microscopy images of droplets containing active WT CASP3 display strong green fluorescence, while droplets containing the inactive D175A variant remain dark. (d) Results of a mock screen demonstrate over 10-fold enrichment of active CASP3.

We tested our emulsion-based assay’s ability to distinguish active CASP3 from an inactive D175A mutant (Fig 1bc). We encapsulated cells expressing each variant and analyzed the droplets using fluorescence microscopy. Droplets that contained active CASP3 displayed a strong fluorescence signal, while droplets with the inactive mutant had no measurable fluorescence. We next tested our assay on-chip using the integrated laser fluorimeter. The active enzyme was easily distinguished from the inactive mutant, with the average CASP3 droplet signal being at least 5 fold greater than the inactive mutant (Supp Fig 1ab).

We next evaluated our microfluidic system’s ability to enrich active caspases from a mixed variant population. We performed a mock sorting experiment by combining active CASP3 with a tenfold excess of an inactive empty plasmid control. We ran this mixed control population through our microfluidic system and sorted droplets with high fluorescence values. We then analyzed the proportion of active CASP3 versus empty plasmid by agarose gel electrophoresis (Supp Fig 1c). We found the initial population contained 9% active CASP3, as expected, and the sorted population contained 95% active CASP3 (Fig 1d). These results indicate that our system can enrich active caspases by at least 10 fold, which is ample for high-throughput screening.

### Deep mutational scanning of the human executioner caspases

Caspases 3, 6, and 7 are referred to as the executioner caspases because they perform the large-scale cellular proteolysis that leads to apoptosis. These enzymes share similar *in vitro* substrate preference, however have been implicated in nonredundant cellular roles that cannot be fully explained by either structural differences or protein expression levels^14–16^. It is likely that subtle differences in their primary amino acid sequence may explain their *in vivo* and *in vitro* functional profiles.

We leveraged our microfluidic screening platform to systematically map sequence-function relationships for CASP3 and CASP7. We generated CASP3 and CASP7 libraries using error-prone PCR. These libraries contained 2-4 amino acid substitutions per variant and approximately 25% of these variants were active caspases. We screened these CASP3 and CASP7 libraries for active caspases using our microfluidic platform. We screened each library in triplicate to evaluate the reproducibility of our methods and to ensure the robustness of our results. For each screening run, we analyzed over 1.5 million caspase variants on average and sorted 4×10^5^ – 7×10^5^ active variants for downstream DNA sequencing analysis (Supp Table 1).

We verified the sorted caspase variants were active enzymes by retransforming the genes into *E. coli* and assaying individual clones in a plate-based format. The initial unsorted libraries were 20-25% functional, while the sorted libraries were 60-90% functional, indicating strong enrichment of functional sequences (Fig 2a). We then sequenced all six sorted samples and their corresponding initial unsorted libraries using Illumina sequencing. The data displayed excellent reproducibility across the three experimental replicates for CASP7 and two experimental replicates for CASP3 (Supp Fig 2). One of the CASP3 replicates did not agree closely with the other two, so we excluded this dataset from the remaining analysis, perhaps due to comparatively poor enrichment increasing the false-positive rate (Fig 2a).

**Figure 2:**
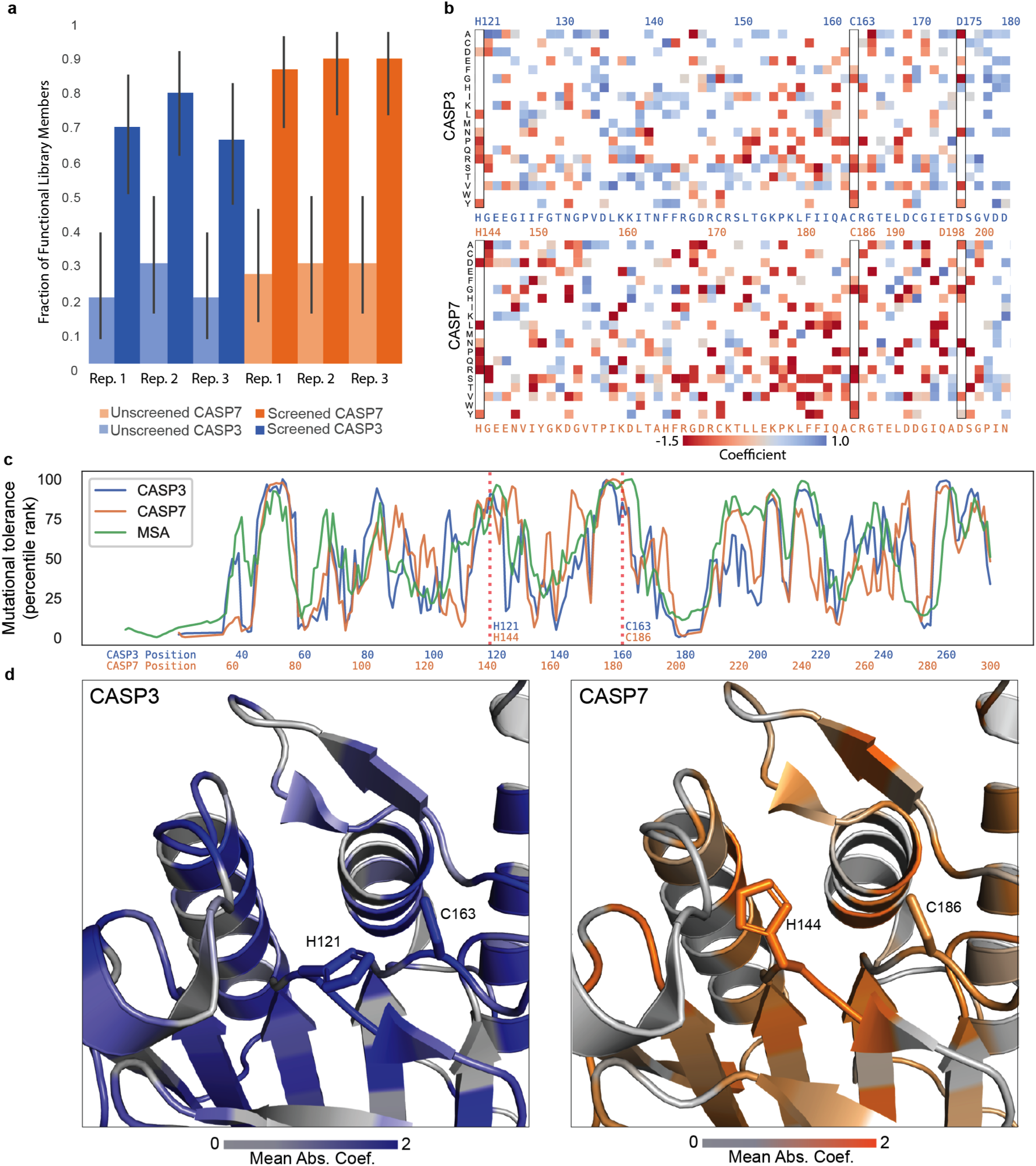
Deep mutational scanning of CASP3 and CASP7. (a) Enrichment of active caspase variants in screened libraries. The fraction of active caspase variants was measured in a plate-based assay before and after screening. The error bars represent the 95% binomial proportion confidence intervals. All replicates showed significant enrichment. (b) A heatmap of mutational coefficients surrounding the active site of CASP3 and CASP7. Mutations that are red have negative coefficients, corresponding to deleterious mutations. Mutations that are blue are positive and are either neutral or activating mutations. White boxes are mutations that did not appear on our DMS analysis. The outlined columns highlight the active site histidine and cysteine residues, as well as the internal aspartate where zymogen maturation occurs. (c) The mutational tolerance of CASP3, CASP7, and the caspase family multiple sequence alignment (MSA) across sequence positions. The mutational tolerance at each position was calculated as the mean absolute value of all mutation coefficients at that position and plotted as a percentile rank. (d) The three-dimensional structures of the CASP3 and CASP7 active sites with their mutational tolerance scores mapped onto the structure. The active site residues are labeled and are strongly colored, indicating low tolerance to mutation.

We used our deep mutational scanning data to build large-scale maps describing how individual mutations affect CASP3 and CASP7 activity (Fig 2b). These maps display expected mutational patterns for both caspases. Substitution of the active site cysteine and histidine residues is highly deleterious. Mutations to large aromatic residues in the hydrophobic core are not tolerated, whereas polar substitutions on the surface of the protein and chemically conservative mutations are generally more lenient. The internal processing sites D175 in CASP3 and D198 in CASP7, which are essential for maturation of zymogenic caspases to mature proteases, are also intolerant to mutation. In addition to corroborating known and expected mutational patterns, our data also revealed new mutations that appear to enhance caspase activity. CASP3 G177R is an activating mutation that is located in an unstructured solvent-exposed loop, where an increased polarity may enhance protein folding and solubility. Another activating mutation, CASP7 F241G, occurs in the hydrophobic core of the protein and may increase the protein flexibility to allow better proteolysis by improving the dynamics of substrate binding to the enzyme active site^17^.

We aggregated the individual mutational effects to obtain the mutational tolerance of each position in CASP3 and CASP7’s primary sequence. This mutational tolerance is related to a site’s importance for caspase function and allows us to analyze broader sequence and structural features. The site-wise mutational tolerance profiles of CASP3 and CASP7 are generally very similar, and also agree closely with profiles generated from a multiple sequence alignment (MSA) of natural caspases (Fig 2c). The beta-sheets that comprise the proteins’ core are less mutable than the exterior helices, and the active site is evolutionarily conserved in the MSA and was also seen to be immutable in our deep mutational scan.

### Contrasting mutational profiles reveals functional differences across executioner caspases

Humans possess 12 separate caspases that all diverged from a common ancestor and share the same structurally conserved proteolytic domain. Despite their highly similar structure and biochemical activity, each caspase’s regulation and cellular targets are unique and confer numerous non-redundant physiological roles. We explored our CASP3 and CASP7 sequence-function profiles to better understand functional differences between highly similar members of the caspase family.

We compared the mutational profiles of CASP3 and CASP7 to identify sites that display differing mutational tolerance and may have functionally diverged during caspase evolution and specialization (Fig 3ab). One notable sequence position was E173 in CASP3 and the equivalent residue Q196 in CASP7 (Fig 3c). CASP3 can tolerate any substitution at this position, whereas CASP7 can only accept substitution to glutamic acid. Intriguingly, Q196 is a known important regulatory site in CASP7 that is cleaved by Cathepsin G to activate procaspase-7^18^. While Cathepsin G is not present in our *E. coli*-based screen, it’s possible that CASP7 can self-activate at this site and amino acid substitutions at this site reduce the pool of active enzyme.

**Figure 3.**
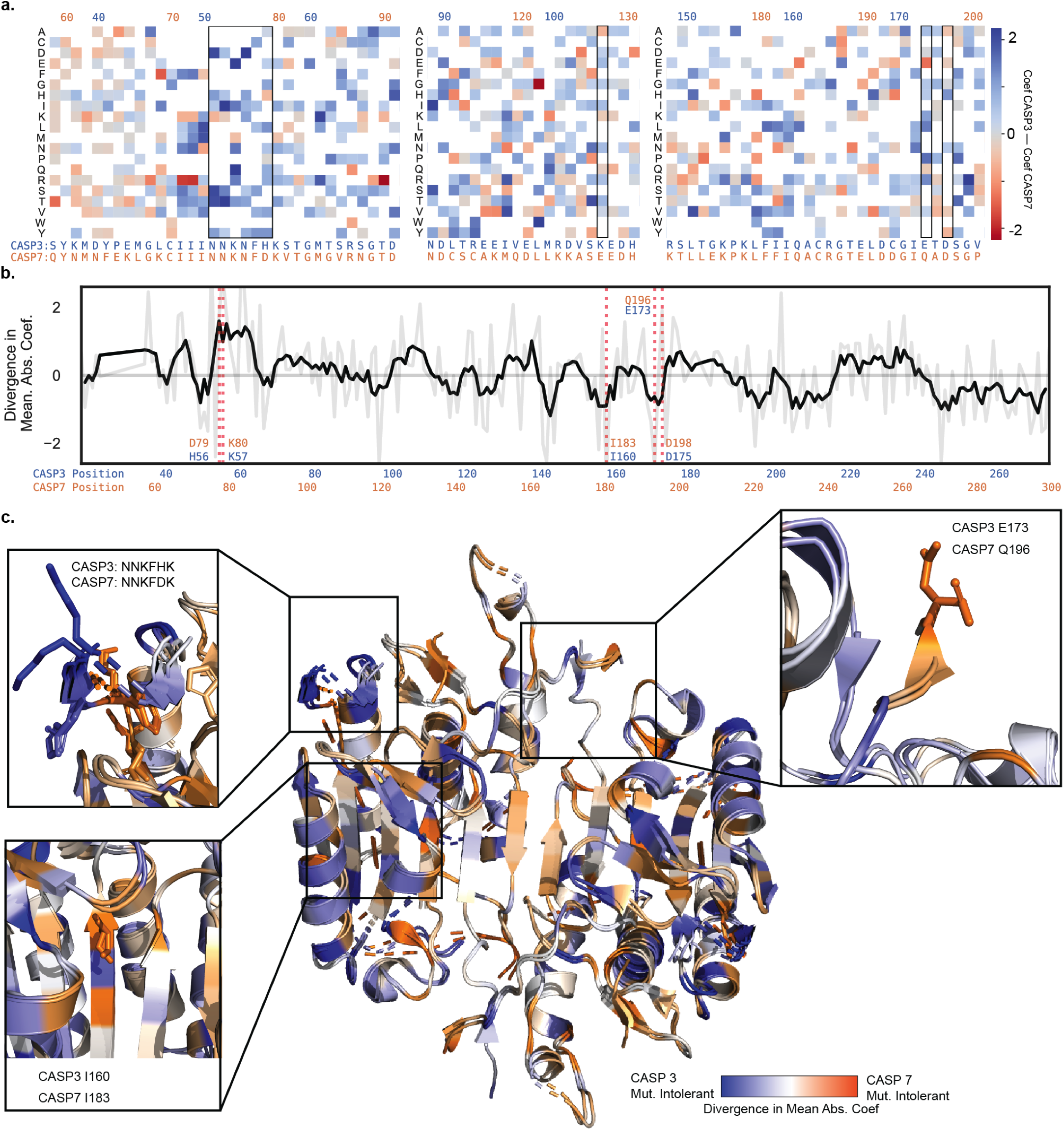
Divergence in Caspase-3 and −7’s mutational landscapes. (a) A heatmap of mutation coefficient differences between CASP3 and CASP7. Blue indicates a larger coefficient in CASP3 and orange indicates CASP7 has a larger coefficient. (b) Differences in mutational tolerance between CASP3 and CASP7. The light grey line shows the difference in a site’s mean absolute coefficient between CASP3 and CASP7, and black line is a moving average to highlight general differences. Positive values are positions where CASP3 has a larger mean absolute coefficient and thus mutations at that site have a larger effect. Negative values are where CASP7 has a larger effect. The red dotted lines indicate positions of interest. (c) A mapping of the difference in mean absolute coefficient onto the aligned CASP3 and CASP7 structures (2H5J and 2QL5, respectively). The expanded boxes highlight the sequence regions shown in panels a and b.

Another differing sequence region were the adjacent sites H56/K57 in CASP3 and the corresponding D79/K80 in CASP7 (Fig 3c). Our analysis indicates that CASP3’s H56 and K57 are completely insensitive to mutation, with the exception of H56P. In CASP7, D79 and K80 show complete mutational intolerance, with all substitutions being deleterious. These residues are located in a solvent exposed loop that displays identical conformations between CASP3 and CASP7 and is located near the substrate binding site. Inspection of the crystal structures reveal that CASP7 has an extensive salt-bridge network in this region, while CASP3 does not. Presumably, mutations in CASP7 disrupt the salt-bridge network and lead to an altered conformation or destabilization in the protein structure.

A final site to note was I160 in CASP3 and I183 in CASP7 that are located within a beta-sheet in the core of the enzyme (Fig 3c). I160T and I160S are well tolerated in CASP3, but CASP7 cannot tolerate any substitutions at I183. The packing environment of I160 and I183 are identical in the crystal structures, with the neighboring sidechains matching with sub-angstrom alignment. It’s possible that substitutions at these sites are always destabilizing, but CASP3 has additional stabilization from elsewhere in the protein to permit these destabilizing core mutations.

## Discussion

Caspases play a key role in numerous biological processes that are important for human health and disease. A deeper understanding of caspase structure, function, and regulation could open the door to novel therapeutic approaches^4^. In this work, we performed deep mutational scanning on CASP3 and CASP7 to reveal differences between highly similar members of the caspase family.

This work was enabled by our high-throughput droplet microfluidic screening platform that analyzes over 300,000 variants per hour in a highly controlled *in vitro* reaction environment. Our device allowed us to have strict control over how long the proteolytic reactions were allowed to occur, which allowed us to more effectively differentiate caspase variants with altered activity. Cell-based assays that rely on proteolytic reporters and fluorescence-activated cell sorting (FACS) occur on much longer timescales and thus cannot distinguish WT-like activity from variants with severely diminished activity. Even catalytically “dead” active site mutants such as CASP3 D175A display enough enzyme activity to hydrolyze all of the substrate within a few hours. The ability to screen caspases based on fast reaction timescales is necessary to distinguish finer functional differences.

We mapped the effects of 1644 amino acid substitutions in CASP3 and 1772 amino acid substitutions in CASP7—roughly one third of all possible single amino acid substitutions. Our results corroborated findings from previous research, such as mutational intolerance of the catalytic cysteine and histidine residues and other known allosteric and processing sites. We also observed mutational constraints in both enzymes that closely follow our understanding of protein stability from the structural perspective, such as the destabilizing effects of disrupting salt bridges or mutations to the hydrophobic cores.

We compared the mutational profiles between CASP3 and CASP7 to identify sites with differing mutational tolerance that may have diverged during caspase evolution and specialization. As expected, a majority of the sites displayed similar mutational tolerance, but a small subset showed statistically significant differences. We identified several key sites that may hold potential for future drug design. CASP7 D79/K80 forms a structurally crucial salt bridge network that is not observed in CASP3. One may imagine designing a drug that could disrupt that network and selectively inhibit CASP7 while leaving CASP3 function relatively untouched.

Our study had several key limitations. First, we chose to express caspases in *E. coli* due to the simplified molecular biology, high transformation efficiency, and the relative insensitivity of bacteria to caspase overexpression. The enzymes expressed in *E. coli* lack glycosylation and must operate in the absence their native regulatory partners such as other caspases and XIAP. In addition, we assayed caspase activity on a single fluorogenic substrate that is likely not fully representative of their diverse cellular targets. These factors could bias our results and reduce the relevance for caspase function in human cells.

Another limitation of deep mutational scanning (DMS) studies in general is the inability to dissect detailed molecular mechanisms. Our DMS measurements describe *how* amino acid substitutions affect caspase activity, but they don’t explain *why*. A mutation that decreases caspase activity could be the result of changes in protein expression, stability and folding, catalytic rate constants, substrate specificity, allosteric regulation, and more. Further biochemical characterization of individual mutants is necessary to obtain a complete picture of inner molecular workings of caspases.

Our results have highlighted a number of interesting future research directions. Residue Q196 appears to play an important role in CASP7 regulation, presumably because it serves as a secondary cleavage site for activation. Previous work found cleavage at the canonical D198 site or Q196 both activate CASP7, however the Q196 isoform is resistant to inhibition by XIAP^18^. The fact that CASP3 does not have a regulatory site analogous to Q196 suggests a novel mechanism to control CASP3 via XIAP, while leaving CASP7 unaffected. Further, exploring the possibility of leveraging sites like CASP7 D79/K80 to develop selective caspase inhibitors could be prudent to the field of drug design. Demonstrating practical translational results from our screen could open up possibilities for using deep mutational scanning for targeted and selective drug design for many other peptide targets.

Developing small molecule modulators that can selectively inhibit or activate members of the caspase family could open the door for novel therapeutics for a wide variety of human diseases. Designing such molecules is incredibly challenging given the highly conserved structures and functions of caspases, and our limited understanding of protein dynamics and regulation. DMS studies could narrow the space of potential target sites by directly and empirically correlating thousands of mutations to their functional effects and finding key protein features that functionally differ in closely related families of proteins.

## Materials and Methods

### Caspase library generation

Δpro-domain *casp3* and *casp7* genes were amplified using error-prone PCR to introduce random mutations. Error-prone PCR was performed following a protocol calling for 50 uM MnCl_2_ to decrease the fidelity of *Taq* polymerase^19^. We did 15 amplification cycles, introducing ~4.5 nucleotide mutations in the gene. We subsequently purified the amplified product, digested it overnight with DpnI to remove remaining wildtype plasmid inserts, and cloned the insert back into *pET22b* using Circular Polymerase Extension Cloning (CPEC)^20,21^.

The CPEC product was purified and used to transform electrocompetent *E. coli* C43(DE3) cells (Lucigen). Transformed cells were recovered for 45 minutes at 37 °C then and diluted into 200 mL of sterile LB media with the added carbenicillin. Once the culture’s optical density (OD) approached the lower detection limit of our spectrometer (OD_600_ = 0.2), the culture was concentrated, and freezer stocks of 25% glycerol were made and stored at −80 °C. Each library had roughly 10^7^ transformants. 10 transformants were picked from each library and their plasmids sequenced to find that each library had ~2.5 amino acid substitutions per library member.

### Plate reader-based caspase activity assay

Individual clones from the mutagenized libraries were incubated in Magic Media (invitrogen) for 18 hours at 30 °C. Cells were pelleted and resuspended in solution 50 mM Tris, ph 7.4, 50 mM KCl after decanting the supernatant media to achieve a density of 1 OD_600_/mL. 200 uL of resuspended culture was added to a black 96-well plate. 200 uL of assay reagent (0.3x BugBuster, 20 uM DEVD-Rhodamine-110, 50 kU/mL Lysozyme, 50 mM Tris pH 7.4, 50 mM KCl, 100 uM EDTA) was added to the plate and the fluorescence (excitation at 480 nm, emission at 530 nm) over time measured on a plate reader. Sequences with >50% of the wildtype activity were considered to be functional.

### Microfluidic device fabrication

An initial layer of photoresist resin, SU-8 3010, is coated onto a mirrored silicon wafer (University Wafers) and spun at 1500 rpm to achieve 15 um layer height. A photomask (Supp Fig 3) of the first layer of the microfluidic device is placed on the layer and 100 J/cm^2^ of UV light is used to polymerize the features. The wafer was baked at 95° C for 10 minutes to catalyze the polymerization. A second 25 um layer of SU8-3025 is coated onto the wafer by spinning at 4000 rpm, and similarly polymerized with the second photomask (Sup Fig 2b) to create the incubation line and baked again. Undeveloped photoresist is washed off with SU-8 developer (1-methoxy-2-propanol acetate, MicroChem).

The wafer is then used to create a relief in un-polymerized PDMS (Dow Corning Sylgard®184, 11:1 polymer:cross-linker ratio), which is then polymerized by baking at 75° C. Inlet and outlet holes are punched with a 0.5 mm biopsy corer. The device is then thoroughly washed with isopropanol and double-deionized water and then plasma treated alongside a clean glass microscope slide, to which it subsequently bonded. Prior to use, microfluidic channels were filled with Aquapel (Pittsburgh Glass Works) to ensure hydrophobicity, and then baked for 10 minutes at 100° C to vaporize any Aquapel left in the channels.

### Microfluidic caspase screening

10 uL of either Caspase-3 or −7 library glycerol stocks was used to inoculate 5 mL of auto-induction media (Invitrogen Magic Media) and allowed to incubate and express for 18 hours at 30 °C. The cultures were pelleted and resuspended in the assay buffer (50 mM Tris pH 7.4, 50 mM KCl, 100 uM EDTA) to a concentration of 0.075 OD_600_ to form the 2x cell suspension. A 2x assay reagent solution of 50 mM Tris pH 7.4, 50 mM KCl, 100 uM EDTA, 0.3x BugBuster, 20 uM DEVD-Rhodamine-110, 50 kU/mL Lysozyme was also made. Both the 2x cell suspension and the 2x assay reagent were loaded into 1 mL luer lock syringes, which were purged of air and fitted with luer-to-PEEK tubing adapters. The cell syringe used PEEK tubing with 0.005” internal diameter, and all other syringes used 0.015” internal diameter PEEK tubing.

Droplets containing expressed Caspase library variants were generated at the co-flow drop maker junction. Both the 2x cell suspension and the 2x assay reagents flowed into the device at 15 uL/hr, and were pinched into droplets by fluorinated oil (HFE 7500) containing 1% (wt/wt) PEG–perfluoropolyether amphiphilic block copolymer surfactant flowing at 100 uL/hr.

After incubating on-chip for ~3 minutes, droplets were sorted using electrocoalescense with an aqueous stream of 10 mM Tris, pH 8, 1 mM EDTA. A 473-nm laser was focused onto the channel just upstream of the sorting junction, each droplet was individually excited, and its fluorescence emission measured using a spectrally filtered PMT at 520 nm. A field-programmable gate array card controlled by custom LabVIEW code analyzed the droplet signal at 200 kHz, and if it detected sufficient fluorescence, a train of seven 180-V, 40-kHz pulses was applied by a high-voltage amplifier. This pulse destabilized the interface between the droplet and the adjacent aqueous stream, causing the droplet to merge with the stream via a thin-film instability, after which the droplet contents were injected into the collection stream via its surface tension. The contents of the sorted droplets were collected in a microcentrifuge tube for further processing. Droplets were processed at 800-1000 Hz. Because the cell occupancy of the droplets was 10%, we analyzed 80-100 cells per second. Caspase-3 and −7 libraries were sorted in triplicate over a total of 6 days. In total we analyzed.

### DNA recovery and sequencing

Recovered plasmid DNA was purified using Zymo spin columns and transformed into ultra-high efficiency 10G Supreme *E. coli* cells (Lucigen). Cells cultured in SOC media and recovered for 45 minutes at 37 °C. The recovered culture was then used in totality to inoculate a larger 200 mL culture which was incubated overnight until its OD_600_ reached 0.5. The larger cultures were pelleted and resuspended in 20 mL 25% glycerol for storage at −80 °C. Dilutions of the culture were plated prior to incubation to measure how many transformants were present. We generally observed 0.75 –1x as many transformants as what we sorted. Plasmid purified from the larger culture was digested with the restriction enzyme DraIII and ScoI, gel extracted, tagmented using the Nextera XT Library Prepration Kit (Illumina) and sequenced using the Illumina MySeq 2×300.

### DMS data processing and analysis

The reads from the Illumina FASTQ files were mapped to the caspase reference gene using Bowtie2^22^, and translated to amino acid sequences. The fitness effect of each observed amino acid substitution was estimated using a positive-unlabeled learning framework that compares sequences from the presorted population with the sorted population^23,24^.

## References

1. Shalini, S., Dorstyn, L., Dawar, S. & Kumar, S. Old, new and emerging functions of caspases. Cell Death Differ. 22, 526–539 (2014).

2. Graham, R. K. et al. Cleavage at the Caspase-6 Site Is Required for Neuronal Dysfunction and Degeneration Due to Mutant Huntingtin. Cell 125, 1179–1191 (2006).

3. Fuentes-Prior, P. & Salvesen, G. S. The protein structures that shape caspase activity, specificity, activation and inhibition. Biochem. J. 384, 201–32 (2004).

4. MacKenzie, S. H., Schipper, J. L. & Clark, A. C. The potential for caspases in drug discovery. Curr. Opin. Drug Discov. Devel. 13, 568–76 (2010).

5. McIlwain, D. R., Berger, T. & Mak, T. W. Caspase functions in cell death and disease. Cold Spring Harb. Perspect. Biol. 5, a008656 (2013).

6. Häcker, H.-G., Sisay, M. T. & Gütschow, M. Allosteric modulation of caspases. Pharmacol. Ther. 132, 180–195 (2011).

7. Deepak, R. N. V. K., Abdullah, A., Talwar, P., Fan, H. & Ravanan, P. Identification of FDA-approved drugs as novel allosteric inhibitors of human executioner caspases. bioRxiv 356956 (2018). doi:10.1101/356956

8. Agniswamy, J., Fang, B. & Weber, I. T. Conformational similarity in the activation of caspase-3 and −7 revealed by the unliganded and inhibited structures of caspase-7. Apoptosis 14, 1135–1144 (2009).

9. Kudelova, J., Fleischmannova, J., Adamova, E. & Matalova, E. Pharmacological caspase inhibitors: research towards therapeutic perspectives. J. Physiol. Pharmacol. 66, 473–82 (2015).

10. Hardy, J. A., Lam, J., Nguyen, J. T., O’Brien, T. & Wells, J. A. Discovery of an allosteric site in the caspases. Proc. Natl. Acad. Sci. U. S. A. 101, 12461–6 (2004).

11. Romero, P. A., Tran, T. M. & Abate, A. R. Dissecting enzyme function with microfluidic-based deep mutational scanning. Proc. Nat. Adad. Sci. 112, 7159–7164 (2015).

12. Fowler, D. M., Stephany, J. J. & Fields, S. Measuring the activity of protein variants on a large scale using deep mutational scanning. Nat. Protoc. 9, 2267–2284 (2014).

13. Fowler, D. M. & Feilds, S. Deep mutational scanning: a new style of protein science. Nat. Methods 11, 801–807 (2014).

14. Brentnall, M., Rodriguez-Menocal, L., De Guevara, R., Cepero, E. & Boise, L. H. Caspase-9, caspase-3 and caspase-7 have distinct roles during intrinsic apoptosis. BMC Cell Biol. 14, 32 (2013).

15. Slee, E. A., Adrain, C. & Martin, S. J. Executioner caspase-3, −6, and −7 perform distinct, non-redundant roles during the demolition phase of apoptosis. J. Biol. Chem. 276, 7320–6 (2001).

16. Lakhani, S. A. et al. Caspases 3 and 7: key mediators of mitochondrial events of apoptosis. Science 311, 847–51 (2006).

17. Fischer, M., Coleman, R. G., Fraser, J. S. & Shoichet, B. K. Incorporation of protein flexibility and conformational energy penalties in docking screens to improve ligand discovery. (2014). doi:10.1038/NCHEM.1954

18. Scott, F. L. et al. XIAP inhibits caspase-3 and −7 using two binding sites: evolutionarily conserved mechanism of IAPs. EMBO J. 24, 645–655 (2005).

19. Bloom, J. D. et al. Evolution favors protein mutational robustness in sufficiently large populations. BMC Biol. 5, 1–21 (2007).

20. Quan, J. & Tian, J. Circular polymerase extension cloning of complex gene libraries and pathways. PLoS One 4, 6441 (2009).

21. Quan, J. & Tian, J. Circular polymerase extension cloning for high-throughput cloning of complex and combinatorial DNA libraries. Nat. Protoc. 6, 242–251 (2011).

22. Langmead, B. & Salzberg, S. L. Fast gapped-read alignment with Bowtie 2. Nat. Methods 9, 357–359 (2012).

23. Song, H. & Raskutti, G. PUlasso: High-Dimensional Variable Selection With Presence-Only Data. J. Am. Stat. Assoc. 115, 334–347 (2020).

24. Song, H., Bremer, B. J., Hinds, E. C., Raskutti, G. & Romero, P. A. Inferring Protein Sequence-Function Relationships with Large-Scale Positive-Unlabeled Learning. Cell Syst. 12, 92–101.e8 (2021).

