## Supplemental information for "Microfluidic deep mutational scanning of the human executioner caspases reveals differences in structure and regulation"

### Table 1

| | Replicate | Drops analyzed | Positive drops sorted | Time (hrs) | Analysis frequency (Hz) | Sorting frequency (Hz) | Fraction of drops sorted | CFU recovered | Fraction functional after sorting |
|---|---|---|---|---|---|---|---|---|---|
| CASP3 | 1 | 15,639,854 | 693,057 | 7.3 | 592 | 26.2 | 4% | 450,000 | 80% |
| | 2 | 6,886,651 | 420,630 | 6.6 | 291 | 17.8 | 6% | 200,000 | 70% |
| | 3 | 15,061,487 | 499,529 | 7 | 598 | 19.8 | 3% | 100,000 | 66% |
| CASP7 | 1 | 19,202,100 | 593,495 | 8 | 670 | 20.7 | 3% | 280,000 | 85% |
| | 2 | 22,474,041 | 602,890 | 8.3 | 755 | 20.3 | 3% | 480,000 | 90% |
| | 3 | 17,601,832 | 414,992 | 7 | 698 | 16.5 | 2% | 120,000 | 90% |

**Supplemental Table 1: Caspase Screening Statistics** CASP3 and CASP7 libraries were sorted in triplicate. Each sorting run lasted for 6.5-8 hours and we were able to recover at least $10^5$ variants per sort.

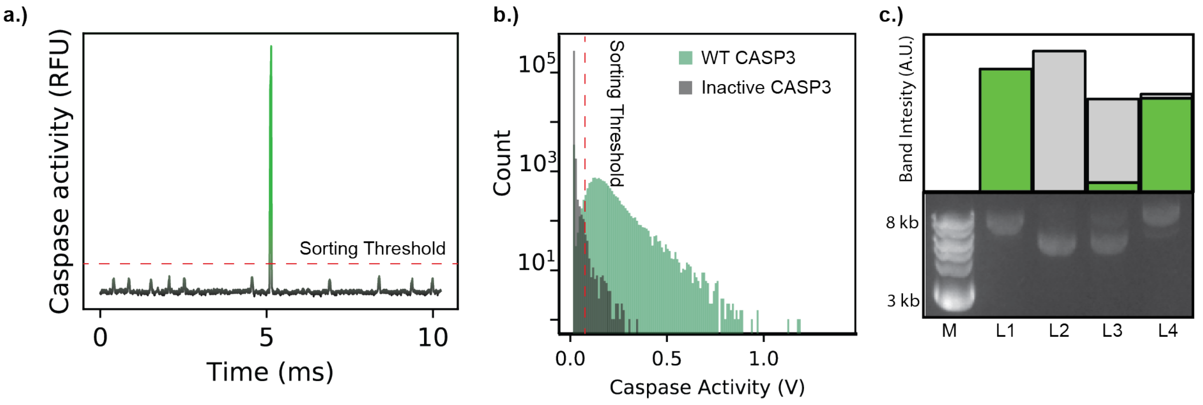


**Supplemental Figure 1:** (a) A time trace of droplets in the microfluidic device as they cross the fluorescence detector. The small peaks correspond to inactive or empty droplets, while the large peak in the center is a droplet containing active CASP3. (b) A histogram of fluorescence activity for WT CASP3 and the inactive CASP3 D175A variant as they are observed in the microfluidic device. WT CASP3 displays significantly higher fluorescence signal than CASP3 D175A droplets. (c) Quantification of recovered plasmid from a mock sorting experiment containing a 10:1 mixture of empty pET22 plasmid to CASP3-containing plasmid. Lanes M: NEB 1kb+ DNA standard; L1: pET 22 plasmid containing WT CASP3; L2: pET 22 plasmid with no insert; L3: a 10:1 mixture of empty vector to CASP3 plasmid containing *E. coli* cells; L4: plasmid recovered after microfluidic screening the L3 input showing significant enrichment of CASP3 plasmid.

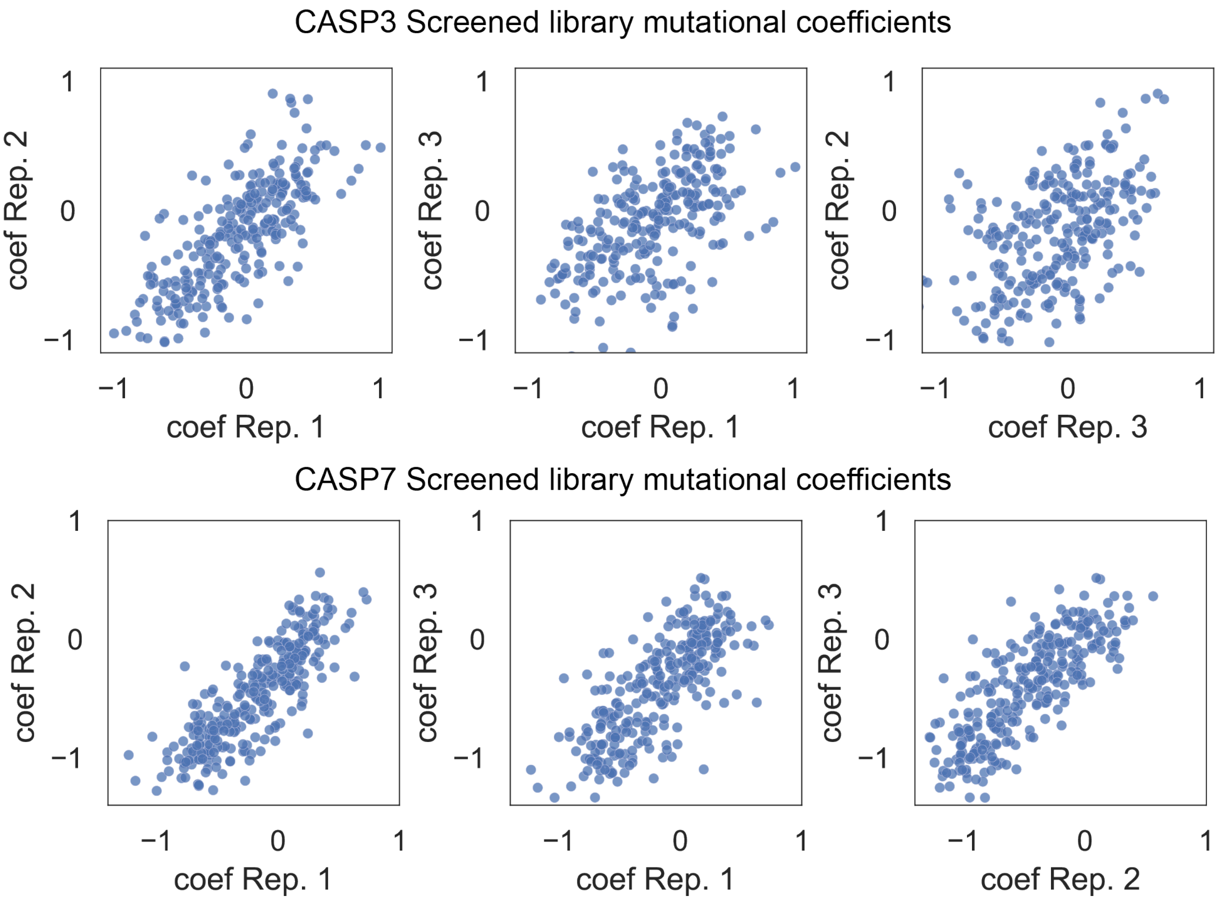


**Supplemental Figure 2:** Correlation of regression coefficients across experimental replicates. For each experimental replicate, the regression coefficient for each mutation is plotted against each other. All three CASP7 experimental replicates correlate well with each other. CASP3 replicates 1 and 2 correlate well with each other, however replicate 3 correlates poorly with the others and was not used for further analysis. It’s possible the microfluidic sorting in replicate 3 had sorting errors that resulted in false positive sequences.

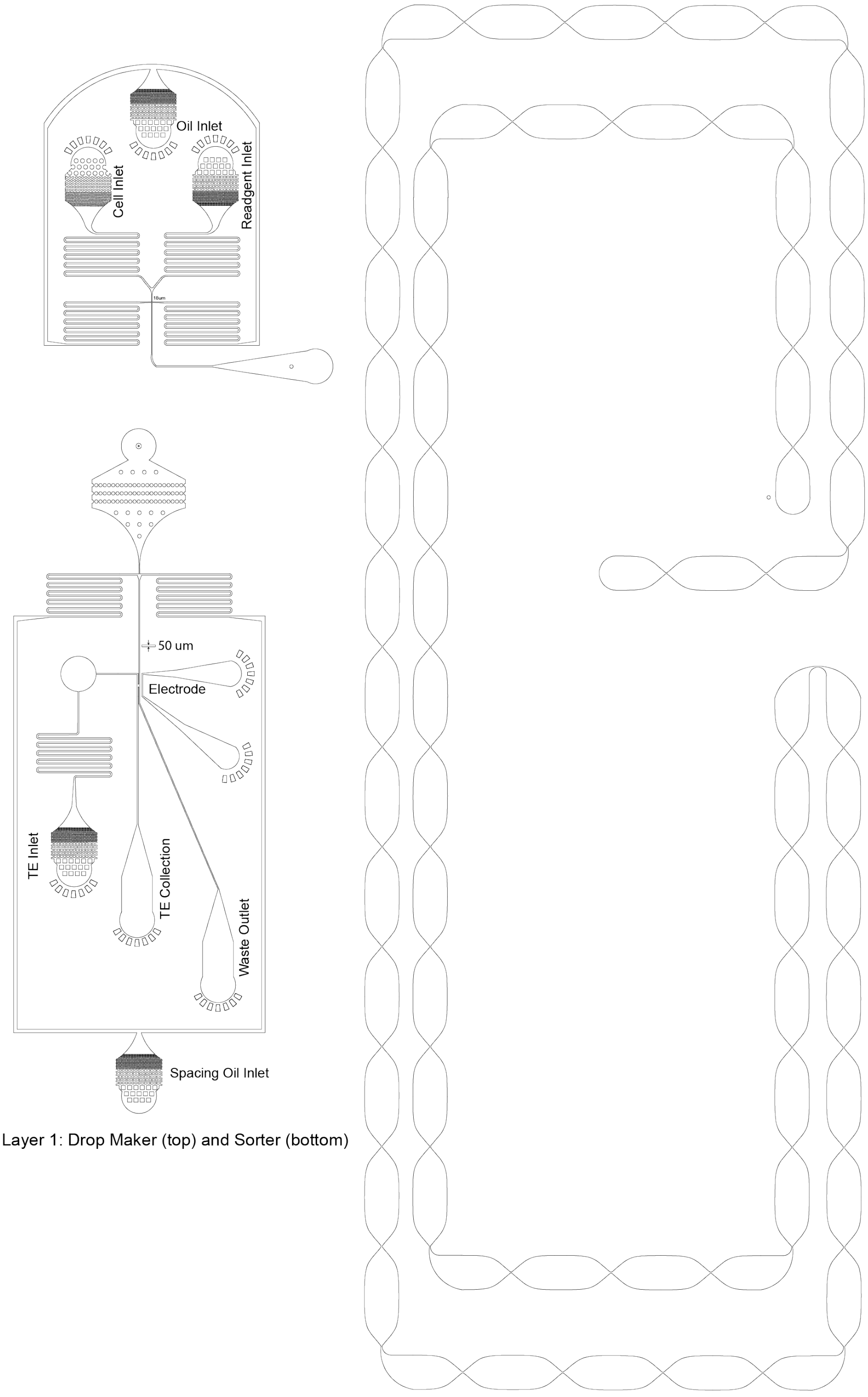


Layer 1: Drop Maker (top) and Sorter (bottom)

Layer 2: Incubation Line

**Supplemental Figure 3:** Microfluidic device designs used in this study. Layer 1 features a 15 um-tall drop-maker and sorter. Layer 2 contains a 50 um-tall incubation line that connects the drop maker and the sorter. The “sausage-like” repeating pattern serves to randomize the position of droplets transverse to the direction of flow to average out the effects of laminar flow that makes droplets in the center move faster than droplets near the edge of the channel.
